# Transverse and axial resolution of femtosecond laser ablation

**DOI:** 10.1101/2022.01.13.476227

**Authors:** Yao L. Wang, Noa W. F. Grooms, Samuel H. Chung

## Abstract

Femtosecond lasers are capable of precise ablation that produce surgical dissections *in vivo*. The transverse and axial resolution of the laser damage inside the bulk are important parameters of ablation. The transverse resolution is routinely quantified, but the axial resolution is more difficult to measure and is less commonly performed. In some *in vivo* samples, fine dissections can also be difficult to visualize, but *in vitro* samples may allow clear imaging. Using a 1040-nm, 400-fs pulsed laser, we performed ablation inside agarose and glass, producing clear and persistent damage spots. Near the ablation threshold of both media, we found that the axial resolution is similar to the transverse resolution. We also ablated neuron cell bodies and fibers in *C. elegans* and demonstrate submicrometer resolution in both the transverse and axial directions, consistent with our results in agarose and glass. Using simple yet rigorous methods, we define the resolution of laser ablation in transparent media along all directions.

## Introduction

The advent of the laser in 1960^1^ was rapidly followed by its application for ablation in a variety of research and medical samples (reviewed in Ref. [2]). Scientists and clinicians recognized that focused laser light can produce highly confined damage to materials, resulting in dissection with resolution unachievable through conventional means. In plasma-mediated laser ablation, the sample absorbs sufficient energy from laser light to vaporize the material^3^. When this absorption occurs on a surface, material is explosively removed. Ablation in the bulk requires laser light to penetrate through the sample to its target. To permit propagation of any appreciable distance, the sample must be relatively transparent to the laser wavelength. Absorption of laser light in the bulk is typically achieved by focusing the laser beam and thereby stimulating nonlinear absorption^4,5^. Depending on the amount of laser energy deposited, the laser generates plasma of various densities, leading to localized damage at the focal volume. At energies significantly above the vaporization energy of the media, an explosion and shockwaves can generate damage beyond the focal volume^6,7^.

The foundational studies on plasma-mediated ablation of water apply to laser ablation of biological media, as they are primarily composed of water. The mechanisms of laser propagation and nonlinear absorption are also similar between many transparent materials. Thus, studies performed on various glasses and transparent semiconductors may be relevant to understanding the processes of laser ablation on biological media. Based on prior studies in ablation of transparent material, key parameters for laser ablation include the intensity thresholds for nonlinear absorption and for ablation. An intensity above the absorption threshold produces strong nonlinear absorption, and energy transfers into the material^4^. Together with the parameters for laser focusing, the ablation threshold sets a focal volume where the laser pulses can vaporize material and create damage.

While assessing the transverse resolution of a laser ablation technique is very common, significantly fewer studies assess the axial, or longitudinal, resolution of ablation. This is partially due to the practical challenges of imaging along the optical axis: laser ablation is often paired with widefield imaging of directions orthogonal to the optical axis. Prior studies have gained optical access to side-views through several methods: In one study, an ablated piece of glass was polished down to access the damage spot^8^. Other studies fractured the glass and then imaged it with brightfield, differential interference contrast microscopy, or scanning electron microscopy following graphite coating^9,10^. Another study used two objectives to image from the side while ablating^11^, and a recent study physically cut a soft sample with a razor blade to expose the damaged spot^12^.

A steadily growing number of studies have characterized laser ablation of biological samples (*e.g*., Refs. [13,14]). Our laboratory studies neurons in the nematode *C. elegans*, in part utilizing femtosecond laser ablation. Given the importance of ablation (and imaging) resolution in the axial direction, we seek to establish rigorous techniques for assessing it. In prior studies we established a technique to measure the illumination resolution of widefield and scanned microscopy^15^, and demonstrated submicrometer transverse resolution of ablation with a similar femtosecond laser^14^. Here, we first measure both the transverse and axial resolution of laser ablation in agarose and glass, materials with permanent visible damage. We find that the transverse and axial diameters of the minimum damage spot are very similar, in both materials. The size of the damage spot over different laser powers obey a simple model of propagation and absorption, from which we can calculate the threshold intensity and the minimum focal spot size^16^. Second, we establish an upper bound on the damage sizes in the transverse and axial directions to define the resolution in *C. elegans in vivo*. Third, we present evidence suggesting that neuron fibers are not damaged by laser irradiation outside of the axial damage region.

## Results

Our setup is shown in Fig. 1a, and a similar setup was described in detail^13^. Various optics and laser parameters impact the laser ablation resolution, including laser wavelength, pulse duration, pulse energy, exposure time, repetition rate, and the numerical aperture (NA) of the microscope objective. In our experiments, most of these parameters are fixed. A femtosecond laser outputs 1040-nm 400-fs pulses at 1 kHz, which we expand and send into a 1.4 NA, 60x objective. In our biological samples, we image epifluorescently through the same objective. We input illumination light into the sample, and we image fluorescent light from the sample onto an array camera.

**Fig 1.**
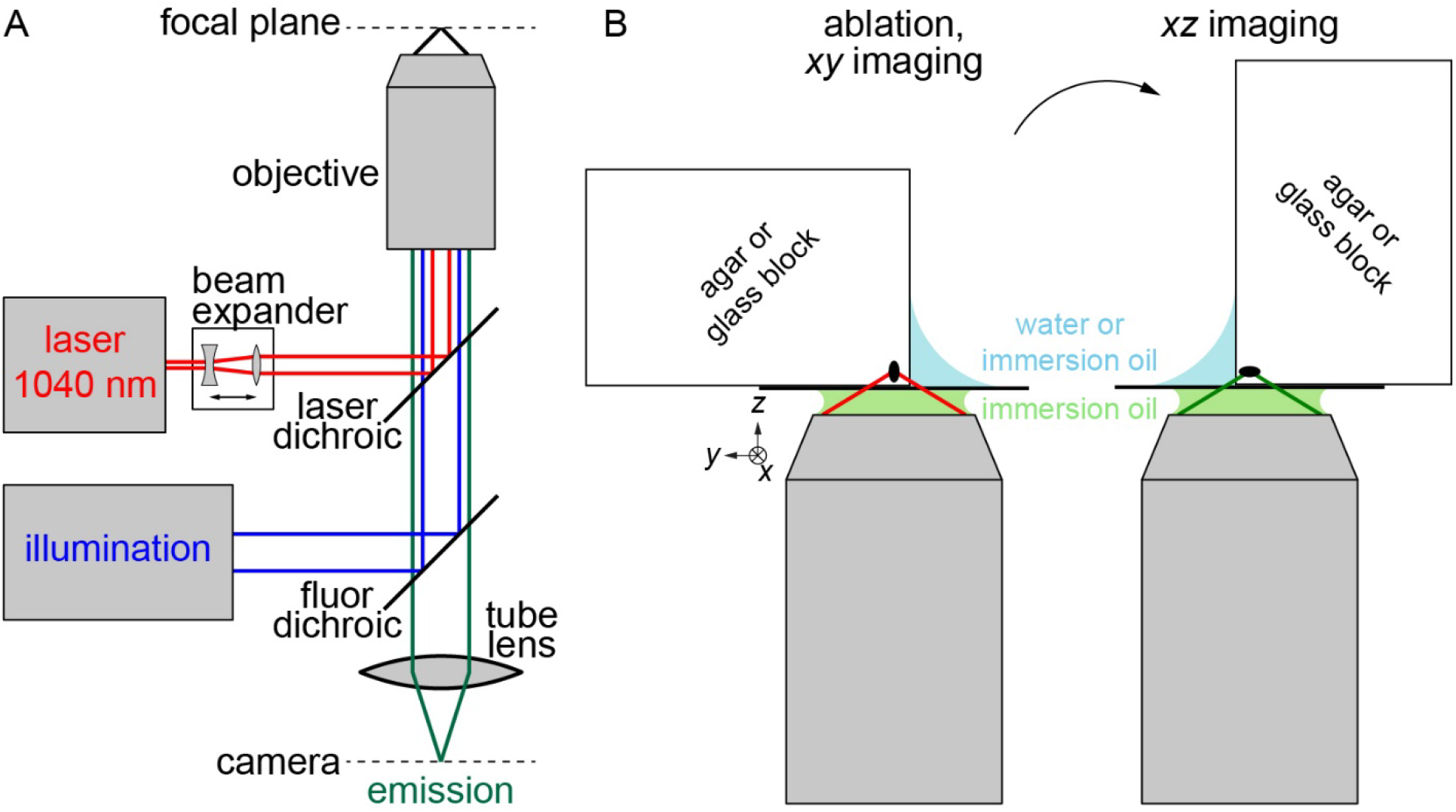
Ablation and imaging setup. (A) Microscope setup for ablation and imaging. 1040-nm femtosecond laser beam is expanded and focused on sample by objective. Illumination light is focused by same objective, and emission light is imaged onto array camera by tube lens. (B) Setup for ablation and imaging. Transverse (*xy*) imaging in same configuration as ablation. Axial (*xz*) plane imaging by ablating block near edge, rotating block 90°, and imaging. Gaps in beampath are filled with water and/or immersion oil to homogenize refractive index.

### Theoretical considerations in a simplified ablation model

This section derives the transverse (*xy*) and axial (*z*) intensity distributions for a focused beam propagating in the *z* direction in a homogeneous medium. We partially follow the derivation in Chapter 2.5 of Ref. [17] with adjustments to follow the derivation in Ref. [16]. We utilize the final results in the following sections to model the ablation we observe and determine the ablation parameters, similar to our prior work that was limited to the transverse direction^14^.

We model focused laser beams with a Gaussian intensity distribution in the transverse direction. Because of diffraction, focused beams come to a minimum width called the waist, at the focal plane (see S1A Fig). The beam width is hyperboloid in the axial direction, and the corresponding intensity distribution is Lorentzian (see S1B Fig). In our simplified model, the laser pulses propagate without scattering, absorption, or nonlinear effects. When the local laser intensity exceeds the intensity threshold for nonlinear absorption, laser energy is deposited into the media. The absorption of sufficient energy generates plasma that vaporizes the material (plasma-mediated ablation). Under our model, ablation occurs when the laser intensity exceeds an threshold for ablation, following similar work examining phase transitions in silicon^16^. In our experience, this simplified model explains many of the key characteristics of the ablation we note in our experiments on transparent materials near the ablation threshold. We remark on the limitations of the model in the discussion.

The local laser intensity is *I*(*x*,*y*,*z*) = *P* / *A* = *P* / *πr^2^*, where *I*, *P*, *A*, and *r* are the optical intensity, optical power, beam cross-sectional area, and beam radius, respectively. Specifically,

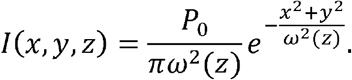

where *P_0_* is the peak power level and 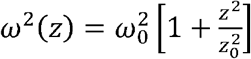 is the squared radius of the beam. *ω_0_* is the beam waist radius at 1/*e*^2^ of the maximum intensity. *z_0_* is the Rayleigh range, or the axial distance from the waist where the intensity drops by half^17^, and 2 *z_0_* is also known as the confocal parameter. Intuitively, the total power at each axial position must be constant if there is no absorption or scattering. Confirming this, the total power is the integral of *I*(*x*,*y*,*z*) over the transverse directions, which yields the peak power, *P_0_*.

We consider simple beams with azimuthal symmetry, so *r*^2^ = *x*^2^ + *y*^2^ and

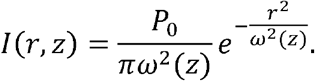

Thus, at the waist (*z* = 0) the intensity is

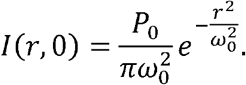

Along the optical axis (*r* = 0) the intensity is

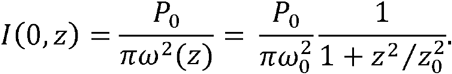

Absorption of laser beams in nominally transparent materials occurs if the laser intensity exceeds a threshold, *I_th_*, dependent on the optical properties of the material^4^. The laser intensity distribution is peaked in the transverse and axial directions. For a given laser power, the intensity meets the threshold at defined transverse *r_th_* or axial *z_th_* radii, where

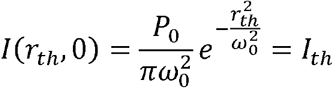

or

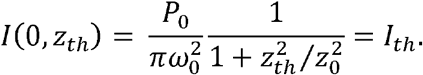

Within this *r_th_* or *z_th_*, laser energy is deposited and material can be ablated, forming a damage spot. The pulse energy, *E*, is the time integral of *P*(*t*), the time evolution of a single pulse’s power. Assuming a Gaussian shape in time, 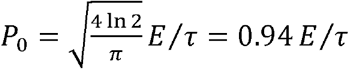, where *τ* = 400 fs is the pulse duration^18^. The average laser power is *P_avg_* = *E* * *f_rep_*, where *f_rep_* = 1 kHz is the repetition rate. Thus, the respective radii of the damage spot in the transverse and axial directions are

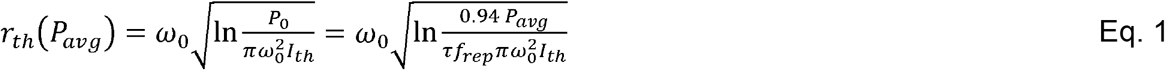

and

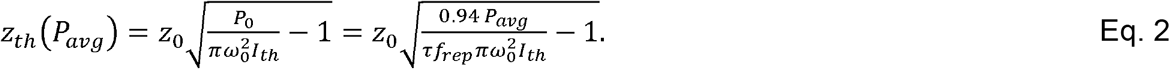

### Agarose and glass ablation

The resolution of laser ablation *in vivo* can be difficult to assess. The media itself is often heterogeneous, complicating observation of laser-induced structural changes. Following ablation of soft tissue, damage locations rapidly flood with water from surrounding regions, which reduces index differences and contrast between the damage spot and the background. We sought to find a biological medium with permanent index changes following ablation to allow precise measurement of damage sizes. We found that ablated agarose has a persistent, albeit small, index change (see Fig. 2ac). Ablation studies have utilized agarose as a proxy for soft tissues due to their similar mechanical and thermal properties^19,20^. For instance, *C. elegans* has a Young’s modulus around 50 kPa, similar to that of agarose gels^21^. The primary component of both agarose and transparent soft tissues is water, and these media have similar refractive indices. The refractive indices of water, 4% agarose, and *C. elegans* tissue are 1.333, 1.339, and 1.379, respectively^22,23^. Because of their similar refractive index, we expect the propagation of light through agarose and *C. elegans* tissue to be similar. While the resolution and threshold of laser ablation in *C. elegans in vivo* is different from agarose, the agarose ablations provide information on the relative size of the axial resolution compared to the transverse resolution.

**Fig 2.**
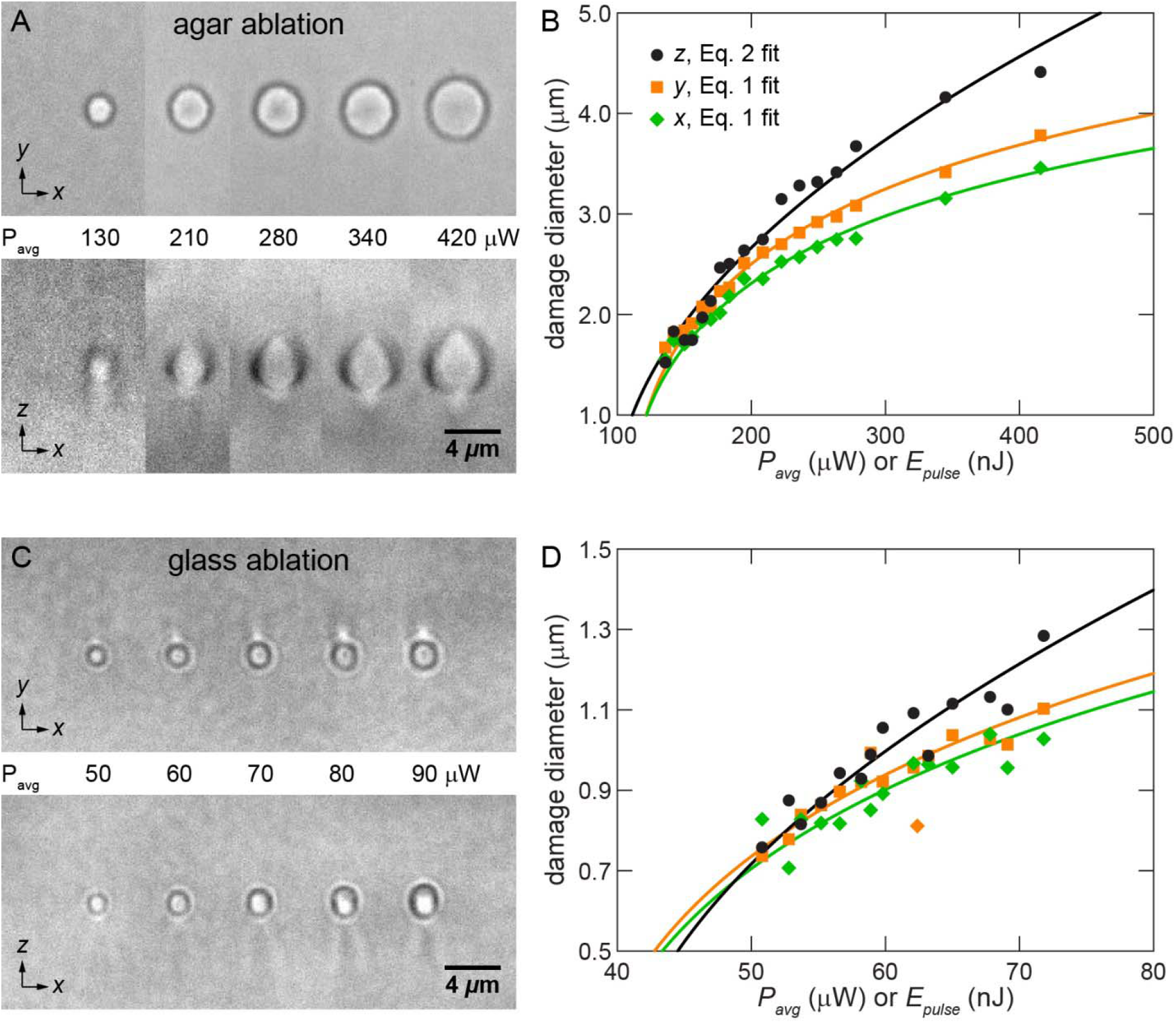
Glass damage spots match theoretical predictions. (A) Transverse and axial plane images of agarose ablation by focused laser pulses. (B) Ablation spot measurements in *x* (diamond), *y* (square), and *z* (circle) directions. Green and orange curves plot Eq. 1 with best fit *ω_0_* and *I_th_*. Black curve plots Eq. 2 with best fit *z_0_* and *I_th_*. (C) Images of glass ablation. (D) Ablation spot measurements in *x*, *y*, and *z* directions, with best fit curves.

Measuring transverse sizes of the damage spot at various pulse energies is relatively straightforward as ablation and reimaging can occur under the same configuration. Measuring the axial size of the damage spot is significantly more difficult. As shown in Fig. 1b, after ablating a block of agarose near a corner, we rotated the block 90° to image the damage spot in the orthogonal (*i.e*., *xz*) direction. We fill spaces between the coverslip and the agarose block with water (blue). The water minimizes changes in refractive index for the portions of the ablation and imaging beams that do not enter the agarose block directly from the coverslip. This homogenization maximizes light transmission and minimizes optical distortion.

We ablated agarose with 100-400 μW average power, producing damage spots of various diameters in *x*, *y*, and *z* (see Fig. 2a). We empirically determined that the agarose damage threshold under our setup is 120 μW average power at 1 kHz repetition rate, or 120 nJ pulse energy. Surprisingly, near the ablation threshold the spot diameters in transverse and axial directions were very similar, producing a roughly spherical damage spot. In the section above, we derive the equations governing the intensity distribution in the transverse (Gaussian) and axial (Lorentzian) directions. As the pulse energy increased, the axial diameter of the damage spot increased more rapidly than the transverse, as expected from Eqs. 1–2. Fitting the transverse agarose damage spot measurements to Eq. 1, we obtain transverse waist radii *ω_0_* in the x and y directions of 1.5 and 1.6 μm, and an ablation intensity thresholds of *I_th_* = 3.7 and 3.1 × 10^12^ W/cm^2^ for ablation of agarose (see Fig. 2b, Tab. 1). We believe the waist radius in the y direction is larger than x direction because some of the beam in the y direction passes through a section of water (see Fig. 1b), which has a slightly different refractive index than agarose. Fitting the axial damage spot measurements to Eq. 2, we obtain an axial radius *z_0_* = 1.3 μm and a separate ablation intensity threshold *I_th_* = 3.3 × 10^12^ W/cm^2^ (see Fig. 2b, Tab. 1). The calculated transverse and axial waist diameters are similar, which is confirmed by roughly spherical damage spots near the threshold.

**Table 1:** Ablation parameters.

|  | Agarose ablation |  |  | Glass ablation |  |  |
| --- | --- | --- | --- | --- | --- | --- |
|  | x | y | z | x | y | z |
| $\omega_0$ ($\mu\text{m}$)<br>$z_0$ ($\mu\text{m}$) | 1.5 | 1.6 | 1.3 | 0.66 | 0.68 | 0.69 |
| $I_{th}$ ($\text{W}/\text{cm}^2$) | $3.7 \times 10^{12}$ | $3.1 \times 10^{12}$ | $3.3 \times 10^{12}$ | $6.5 \times 10^{12}$ | $6.0 \times 10^{12}$ | $6.5 \times 10^{12}$ |

We obtained similar results with the same trends by ablating a thin cylinder of agarose in a water-filled channel (data not shown). Compared to the agarose block, the ablation resolution in the cylinder was not as fine and the imaging was not as clear. We speculate that this is due to refraction from the curvature of the cylinder and increased depth of ablation and imaging. We did not pursue ablations in this configuration further.

We carried out ablations in N-BK7 glass to confirm the similarity between the axial and transverse resolution that we observed in agarose. Ablation of glass leads to a permanent and distinct change in the refractive index that allows a more precise measurement of the damage spot than is possible in agarose. We also ablate glass with power levels that are closer to levels used in ablations of *in vivo* samples. We followed a similar procedure to the agarose ablations above but used immersion oil around the block (see Fig. 1b). We ablated glass with 50-90 μW (see Fig. 2b) and found that the ablation threshold is 50 μW average power, or 50 nJ pulse energy. Results from ablation of glass confirmed the overall pattern of results in agarose (see Fig. 2b, Tab. 1). Transverse and axial sizes of damage spots near the threshold were very similar and the smallest spots appeared spherical. The calculated waist radii *ω_0_* and confocal parameter *z_0_* were also similar. The ablation intensity threshold was about *I_th_* = 6.4 × 10^12^ W/cm^2^.

### *In vivo* ablation

We also characterized the transverse and axial resolution of our surgeries *in vivo* on neurons in *C. elegans*. Following our prior studies^13,14^, we first assessed transverse resolution by ablating the middle dendrite in a tight bundle of dendrites. We ascertained true ablation rather than photobleaching as the diffusible green fluorescent protein (GFP) label does not return to the ablated region following irradiation^13^. The first ablation in Fig. 3a (after #1) shows a cut of resolution better than 1.03 μm. The second ablation (after #2) shows a cut of resolution better than 0.75 μm. The first cut widens between the “after #1” and the “after #2” images due to dendrite tension, further confirming a complete cut. These data indicate an *in vivo* transverse resolution of well below 1 μm, a similar resolution to our prior work using 800-nm Ti:sapphire and 1030-nm Yb-doped fiber lasers^13,14^.

**Fig 3.**
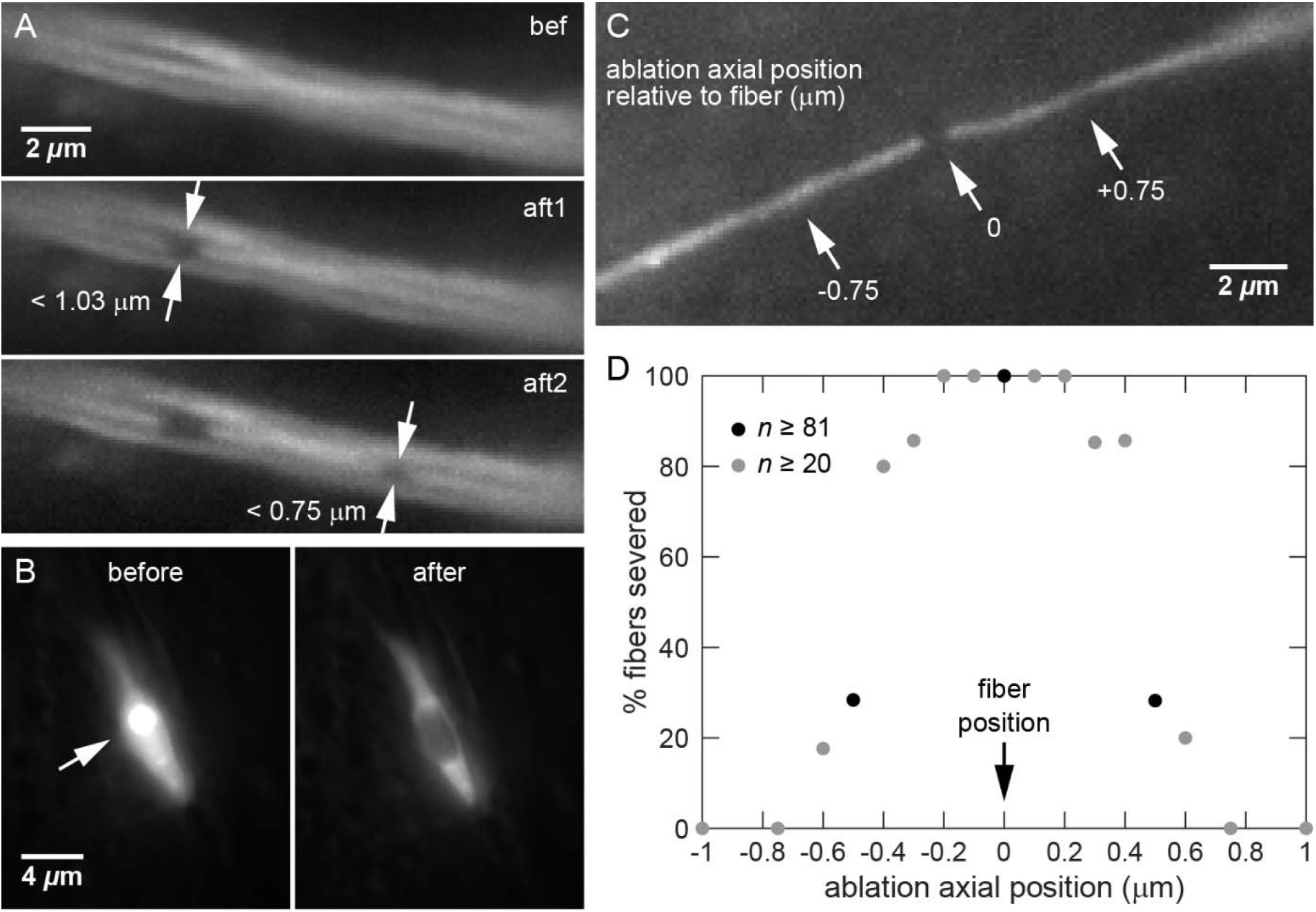
Characterization of *in vivo* laser ablation. (A) Individual neuronal fibers in tight bundle before, after 1^st^ ablation, after 2^nd^ ablation. Images indicate submicrometer transverse resolution of laser ablation. (B) Ablation of nucleosol but not cytosol shows persistence of nuclear membrane, suggesting axial resolution well below 2 μm. “After” image taken immediately after ablation. (C) Fiber severed by ablation directly at its axial position but spared by ablation 0.75 μm above or below fiber. (D) Percentage of fibers severed by ablation at various axial positions relative to the fiber. Results indicate submicrometer axial resolution of laser ablation.

Second, in multiple studies, we have ablated cell bodies for several seconds to kill off entire cells^13,24^. We visualize and target single cells by a diffusible but cell-specific GFP label. GFP diffusion through the fluid cell cytosol and nucleosol is very rapid, but its diffusion across membranes in *C. elegans* neurons is much slower, on the order of tens of seconds. In the early stages of cell body ablation, we often find that the nucleus dims while the cytosol remains fluorescent (see Fig. 3b). This dimming occurs because the interior of the nucleus is ablated while the nuclear membrane is spared. GFP from throughout the nucleosol diffuses to the laser focus and is selectively ablated, decreasing the fluorescence of the entire nucleus. The intact nuclear membrane greatly slows the diffusion of cytoplasmic GFP into the nucleus and preserves the fluorescence of the cytosol. As shown in Fig. 3b, irradiating the bright nucleolus (arrow) leads to dimming of the entire nucleus but not the cell cytosol, confirming that ablation does not damage the nuclear membrane. Thus, the axial resolution of laser ablation is far below 2 μm, the axial depth of the round nucleus.

Third, to further define the axial resolution, we focused laser pulses at specific locations below, directly on, or above single neuronal fibers. As shown in Fig. 3c, ablating at the fiber’s axial position severs the fiber while ablating a sufficient distance above or below the fiber does not sever the fiber. We tracked the percentage of fibers severed by focusing at each depth relative to the fiber’s axial position. As shown in Fig. 3d, the efficacy of laser ablation peaks when the laser pulses are focused on or near (≤ 0.2 μm) the fiber. Surgical efficacy decreases roughly symmetrically as the focus moves away from the fiber axially. At approximately ± 0.4-0.5 μm there is a clear transition from severing to non-severing. Laser pulses focused at 0.75 μm away from the fiber or further are unable to sever the fiber. There is noticeable pulse-to-pulse variability in the laser’s operation, producing a non-binary distribution. Sequential pulses can focus on slightly different transverse and axial positions over a range of ~0.1-0.2 μm. Thus, these data indicate that the axial resolution of laser ablation is slightly below 1 μm.

Lastly, we cut neurites in the ASJ neuron and noted axon regeneration after ablation to provide another demonstration of surgical resolution. As shown in Fig. 4a, the ASJ neuron consists of a cell body (light gray), dendrite (black), and axon (dark gray). In mutant animals with defective dual-leucine zipper kinase, denoted as *dlk-1*, multiple neuron types have severely reduced axon regeneration^25^. The ASJ axon shows no regeneration in *dlk-1* animals when it is the sole fiber cut; however, nearly 100% of ASJ axons regenerate when the ipsilateral sensory dendrite is severed concomitantly^26^. In *dlk-1* animals, we cut the ASJ axon by ablating directly on the axon and then immediately ablated at various depths relative to the dendrite. As shown in Fig. 4b, *dlk-1* animals with axon cuts but no dendrite ablation showed no regeneration, confirming prior results. When we cut axons and ablate directly on the dendrite, 95 ± 5% of neurons regenerate. When the dendrite is ablated 0.5 μm above or below its depth without severing it, there is minimal regeneration. These data further confirm the tight confinement of laser damage to the focal plane.

**Fig 4.**
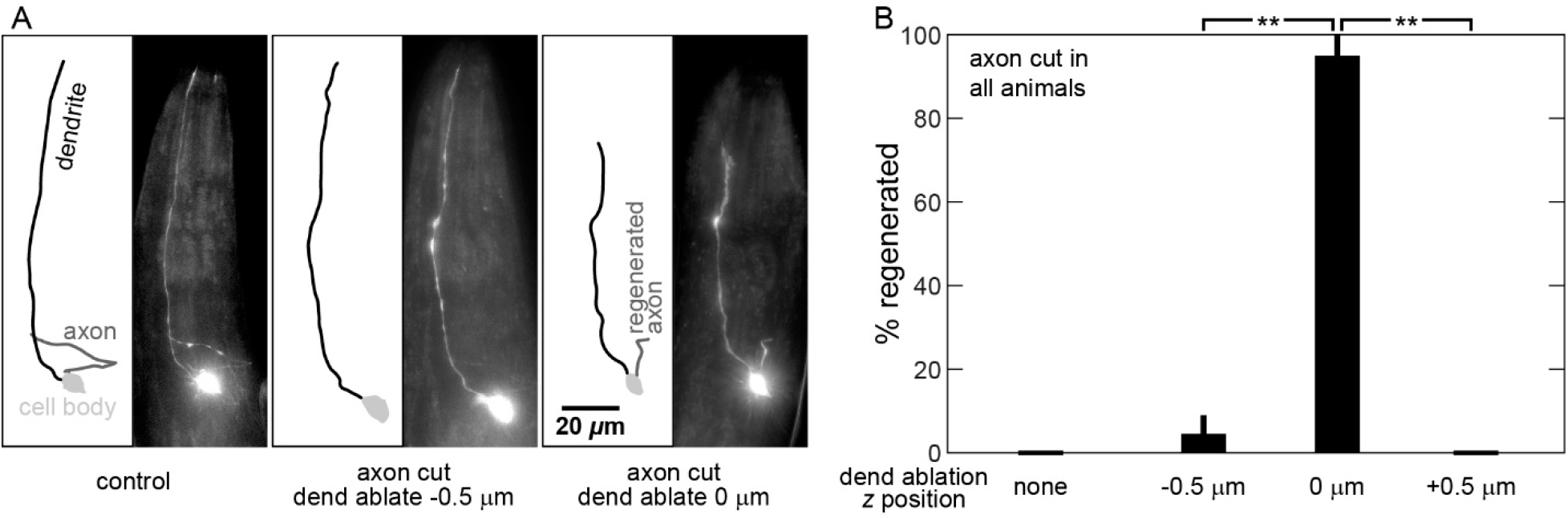
Regeneration assay confirms submicrometer resolution of laser ablation. (A) Line drawings and fluorescent images of ASJ neuron. Control (left) neuron with cell body (light gray), intact dendrite (black), and intact axon (dark gray). Non-control images taken 48 hours after ablation: Postablation neuron (middle) with cut axon and ablation 0.5 μm below dendrite. Axon has decayed away. Postablation neuron (right) with regenerated axon and dendrite ablated directly at its axial position. (B) Percentage of neurons regenerating following axon cut only (none) or axon cut with dendrite ablation at various depths. Significant regeneration only occurs when dendrites are ablated directly at their axial position. All animals are *dlk-1* mutants with axons cut. Average and standard deviation indicated by bars. ** *p* < 0.0001.

## Discussion

While both the transverse and axial resolution are needed to fully characterize laser ablation, the axial resolution is rarely measured. Here we determine the transverse and axial resolution of a Yb-doped diode-pumped solid-state femtosecond laser by ablating transparent materials and *in vivo* biological samples. By solving the Gaussian and Lorentzian intensity distributions for *r_th_* and *z_th_*, we find that the transverse diameter of ablation is proportional to the square root of the logarithm of *P_avg_*, while the axial diameter of ablation is proportional to the square root of *P_avg_*. By ablating agarose and glass at various powers, directly measuring the damage spot in all three dimensions, and fitting the measurements to the equations derived, we find an intensity threshold of 3.4 × 10^12^ W/cm^2^ for agarose ablation and 6.4 × 10^12^ W/cm^2^ for glass ablation. Fitting our transverse and axial data led to similar results in all three dimensions for damage size and ablation threshold. Our results on glass are roughly consistent with prior studies showing intensity thresholds of 8 - 20 × 10^12^ W/cm^2^ of various glasses under femtosecond pulses of different wavelengths and pulse durations (*e.g*., Refs. [27,28]). Importantly, we demonstrate, in agarose and in glass, that the transverse and axial resolution of laser ablation near the ablation threshold are very similar to each other (see Fig. 2bd).

Our measurements of laser ablation spots in transparent materials obey our model remarkably well, despite our model’s simplified approach. Clearly, there are many other effects involved in femtosecond laser propagation, absorption, and ablation: First, there are several nonlinear propagation effects, including self-phase modulation, self-focusing, and self-steepening^29^. We believe these effects are minimal because tight focusing conditions minimizes the distance over which these nonlinear effects accumulate. However, we note that we have not precompensated (*i.e*., prechirped) for chromatic dispersion of the beam as it travels from the laser to our sample. By precompensating, it may be possible to reduce the energy needed to achieve the ablation threshold, which could improve ablation resolution. Second, there is likely a small amount of scattering or linear absorption throughout the beam, as deeper targets require more power to ablate. Moreover, the threshold condition for nonlinear absorption is an approximation for determining where absorption occurs. In reality, the absorption is proportional to the intensity raised to the order of the nonlinear absorption process^4^. Third, the absorption mechanisms that generate plasma are much more complicated than the simple nonlinear absorption and photoionization of our model. In fact, avalanche ionization is the mechanism that absorbs the most laser light^6^. However, nonlinear absorption dictates where the absorption occurs and thus the spatial extent of the plasma-mediated damage. Finally, laser repetition rate impacts damage mechanisms, particularly at higher rates, when laser pulses deposit energy faster than it can diffuse away as heat, leading to cumulative thermal damage. Our laser operates well below the 1 MHz repetition rate boundary for cumulative damage^30,31^. Even so, we have seen some cumulative damage in agarose at power levels above those in Fig. 2b, as evidenced by divergence between single pulse and multi-pulse ablation (data not shown).

We also characterized the resolution of 1040-nm femtosecond laser ablation in *C. elegans* neurons *in vivo*. Similar to our prior studies utilizing 800-nm pulses^13^, we show a transverse resolution that is submicrometer by ablation of tight fiber bundles. Likewise, we show an axial resolution that is also submicrometer by observing the frequency of successful severing and surgical impact on regeneration. Note that even though the laser pulses pass through the neuronal fibers (Fig. 3c), the fibers are not damaged unless the region where the laser intensity exceeds the threshold impinges on the fiber. It is only in this region that laser light is absorbed and damage occurs. Fundamentally, it is this nonlinear absorption that confines axial damage so tightly to the focal volume and ultimately produces the high axial resolution.

An increasing number of studies utilize laser ablation in biological contexts. Laser ablation allows selective submicrometer-resolution damage in the bulk at a user-defined timepoint, which is unachievable by conventional methods such as genetics, pharmacology, or manual dissection. Rigorously defining ablation resolution in all three dimensions is crucial for understanding its capacity to produce highly precise dissections without collateral damage. To our knowledge, our study is one of relatively few studies that examine the axial resolution of plasma-mediated laser ablation in transparent media or biological samples. In our *in vivo* experiments, defining the axial resolution is all the more important because the epifluorescence imaging that guides laser ablation has lower resolution in the axial than transverse directions. Ascertaining that the resolution of laser ablation is similar in all three dimensions ensures high quality dissections that are independent of the sample orientation.

## Materials and methods

### Microscope setup

We utilized a Nikon Ti2-E inverted microscope with a SOLA SE II LED light engine to take 3D image stacks. We used a 1.4 NA, 60x oil-immersion objective for ablation and imaging. This objective is designed for use with a #1.5 glass coverslip.

### Laser setup

A Yb-doped diode-pumped solid-state laser (SpectraPhysics Spirit-1040-4W) outputs 1040-nm 400-fs pulses with a repetition rate of 1 kHz. As shown in Fig. 1a, we expanded the diameter of the laser beam through a 10x beam expander (Thorlabs GBE10-B) to overfill the back aperture of our microscope objective. We raised the beam using a custom periscope (not shown) to send into the upper turret of our epifluorescence inverted microscope. A mechanical shutter in the upper turret controls the laser irradiation. We illuminated with blue light for imaging through the lower turret, and the emission light was imaged by an sCMOS camera (Andor Zyla 4.2).

### Agar ablation and imaging

Our ablation and imaging setup is shown in Fig. 1b. We utilized a 4% agarose block (Fisher BioReagents BP160-500). To fabricate the block, we poured molten 4% agarose solution into an open box made from glass slides. After solidifying, we cut the block and immediately immersed it in water to prevent its sharp edge from drying out and deforming. We placed this block onto a coverslip typically used for imaging with our objective. We added pure water to all locations that transmit the laser beam or illumination light. This water prevents impinging of illumination or laser light on a low-index region of air that could lead to refraction or reflection rather than transmission. We ablated agarose 40 μm below the surface with a 0.2 s exposure time. We acquired the damage spot *xy* images by brightfield imaging through the same face as laser transmission. For *xz* imaging of the laser damage spot, we separated the agarose block from the coverslip after ablation, rotated it by 90°, and repeated the same process of attaching the coverslip and imaging on the orthogonal face.

### Glass ablation and imaging

The glass ablation procedure (see Fig. 1b) was very similar to the agarose ablation procedure with the following alterations: We performed all glass ablation on an N-BK7 block of glass polished to 20-10 scratch-dig. We added immersion oil to all the locations that transmit the laser beam or illumination light. We carried out ablation and imaging with procedures similar to the agarose imaging above.

### Damage spot characterization

Ablation of agarose or glass creates a damage spot that refracts light in brightfield images. At the focal plane, the interior of the spot is bright. In Fiji/ImageJ (https://imagej.net/software/fiji/), we measured the size of a damage spot using the intensity profile of a line through the center of the spot. We defined the size to be the inner diameter of the spot, where the intensity profile intersects the background value. We utilized these sizes together with the average laser power to fit Eqs. 1–2 and determine the minimum spot *ω_0_*, *z_0_*, and the intensity threshold via custom MATLAB code.

Measurements in the *y* direction were consistently larger than in the *x* direction, especially in agarose (see Fig. 2bd). We believe this is due to a portion of the beam in the y direction impinging on different media (water instead of agarose, or immersion oil instead of glass, see Fig. 1b). The slight index mismatch may reduce the resolution of the damage spot or its imaging. In agarose calculations, we set *ω_0_* in Eq. 2 equal to the calculated ω_0_ in the *x* direction (see Tab. 1). In glass calculations, the refractive indices and calculated ω_0_ are very similar, thus, we set ω_0_ in Eq. 2 equal to the average of the calculated ω_0_.

### *C. elegans* cultivation, immobilization, imaging

We followed established procedure for *C. elegans* strains cultivation on agar plates^32^ at 15 or 20 °C, animal immobilization by sodium azide, and imaging^13^. After immobilization, animals were rotated to a desired orientation^4^ under a fluorescence stereomicroscope and then imaged under an inverted microscope.

### *C. elegans* strains

We used the following strains for this study: *dlk-1(ju476)I; ofIs1[lin-15ab*+*; trx-1::gfp]IV* (for regeneration ablations), *tax-2(p691)I; ofIs1[lin-15ab*+*; trx-1::gfp]IV* (for nuclear and dendrite ablations), and NG3146 *gmIs18[ceh-23::gfp; rol-6]X* (for dendrite bundle ablations).

### *In vivo* ablation

We followed preoperative procedures established for laser ablation in *C. elegans*^13,33^. Animals carried fluorescently-labeled neurons. We mounted young adults onto agarose pads with sodium azide for immobilization. We followed established procedures to sever neurites by ablating them with 35-nJ pulses for 0.2 s ^4,13^. Neurites are approximately 10 μm deep in the animal. To characterize the transverse resolution of laser damage, we severed tightly-bundled fibers in the amphid dendrite bundle (Fig. 3a). We measured the gap between unsevered fibers using ImageJ. For characterizing axial resolution, we aimed our laser at the ASJ nucleolus, which is 10-20 μm deep, and ablated it with 50-nJ pulses for 15 s (Fig. 3b), following established protocol^13^. We also ablated at defined locations above or below the dendrite (Fig. 3cd) and noted percentage of successful severing.

### Regeneration characterization

We followed procedures previously established in our laboratory for studying regeneration^26^. In young adult animals we severed the axon, the axon and ipsilateral dendrite, or the axon and then ablated 0.5 μm above or below the ipsilateral dendrite without severing (Fig. 4ab). We followed postoperative procedures to recover worms in nematode growth medium and continued cultivation at 20°C. We reimaged worms 48 hours post-ablation and noted the frequency of regeneration.

### Statistics and interpretation of results

We calculated *p*-values for frequency of successful cuts and regeneration with Fisher’s exact test.

## Acknowledgements

We thank the members of the Chung Laboratory, including S. C. Civale and N. Patel, for feedback on the manuscript. NG3146 is a gift from Gian Garriga.

---

**S1 Fig.**
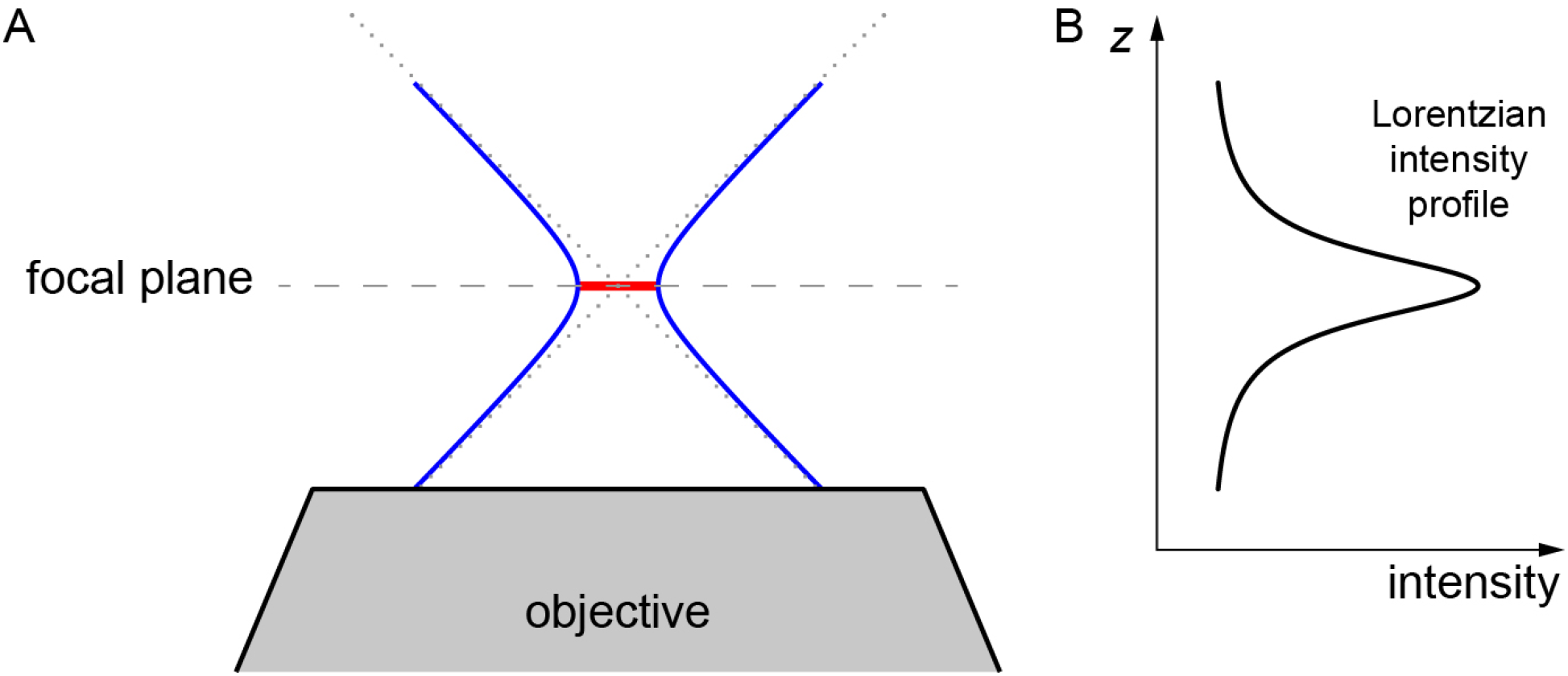
Focused beam profiles. (A) Focused beams are hyperboloid in the axial direction. Asymptotes indicated by dotted lines. Waist diameter indicated by red line on focal plane (dashed line). (B) Corresponding intensity profile of beam shown in (A).

